# Assessment of patient-specific efficacy of chemo- and targeted-therapies: a micropharmacology approach

**DOI:** 10.1101/236653

**Authors:** Aleksandra Karolak, Branton Huffstutler, Zain Khan, Katarzyna A. Rejniak

## Abstract

Both targeted and standard chemotherapy drugs are subject to various intratumoral barriers that impede their effectiveness. The tortuous vasculature, dense and fibrous extracellular matrix, irregular cellular architecture, and nonuniform expression of cell membrane receptors hinder drug molecule transport and perturb its cellular uptake. In addition, tumor microenvironments undergo dynamic spatio-temporal changes during tumor progression and treatment, which can also obstruct drug efficacy. To examine these aspects of drug delivery on a cell-to-tissue scale (single-cell pharmacology), we developed the *microPKPD* models and coupled them with patient-specific data to test personalized treatments.

## I. Introduction to single-cell cancer pharmacology

The clinical success or failure of targeted and chemotherapy treatments depends on how well a drug’s molecules reach all tumor cells (pharmacokinetics, PK) and engage with their molecular targets to invoke the desired therapeutic effect (pharmacodynamics, PD). Conventional PK/PD analyses assess treatment efficacy of the entire organ level, however how drugs penetrate *in vivo* tissues and how they interact with tumor cells are still poorly understood. Recent advancements in imaging techniques have given rise to a new research area of *in vivo* single-cell pharmacology [1-3]. The *microPKPD* (micro-scale PK/PD) modeling framework brings additional power for high throughput simultaneous examination of various physicochemical properties of both drugs and tissues, which provides a tool for improving both drug design and administration protocols that aim to increase anti-cancer treatment efficacy.

## II. Illustrative Results of *microPKPD* Applications

The crucial advantage of the *microPKPD* model is the use of digitized tumor tissues as the model domain. This allows for the testing of the treatments that are calibrated to a specific tissue structure, a particular membrane receptor expression, and precise drug-tumor cell interactions. Here, we discuss the application of *microPKPD* to both biological and medical data on different scales, including patients’ biopsies, tumor organoids, and tumor tissue slices.

### A. Predicting a patient’s tumor chemoresistance with the Virtual Clinical Trials concept

The *Virtual Clinical Trials* concept [4] uses standard-of-care (SOC) medical histology images that are routinely collected in the clinic, advanced image analysis algorithms, and computational simulations to determine whether a patient’s tumor is resistant to SOC treatments. We envision that the patient will follow the schematics presented in Figure 1. Tissue samples will be collected from a routine biopsy procedure (A). Following current clinical practices, the tissue will be stained with hematoxylin and eosin (H&E), sliced, fixed on a glass slide (B), and examined by a pathologist (C). For virtual trials analysis, it will also be scanned and digitized (D). The magnified, high-resolution images will be used to identify and quantify the immunohistochemical (IHC) and morphological features of individual cells (D2, Pathomics). These will be used to characterize the tumor’s metabolic landscape pathology [5,6] and to define cellular phenotypes (E, Virtual Pathology). The patient’s digitized tissue slices (the z-stack) will be used to reconstruct the entire 3D tumor organoid (vasculature and cellularity), and the *microPKPD* model will simulate the drug(s) penetration and tumor response curves (E2). The likelihood of the tumor being either responsive or resistant to the therapy will be calculated to support clinical decisions (F). In addition, the virtual clinical trials model can be used to predict optimal drug administration protocols (the order, dosage, and timing of a combination of drugs) for personalized treatment.

**Figure 1.**
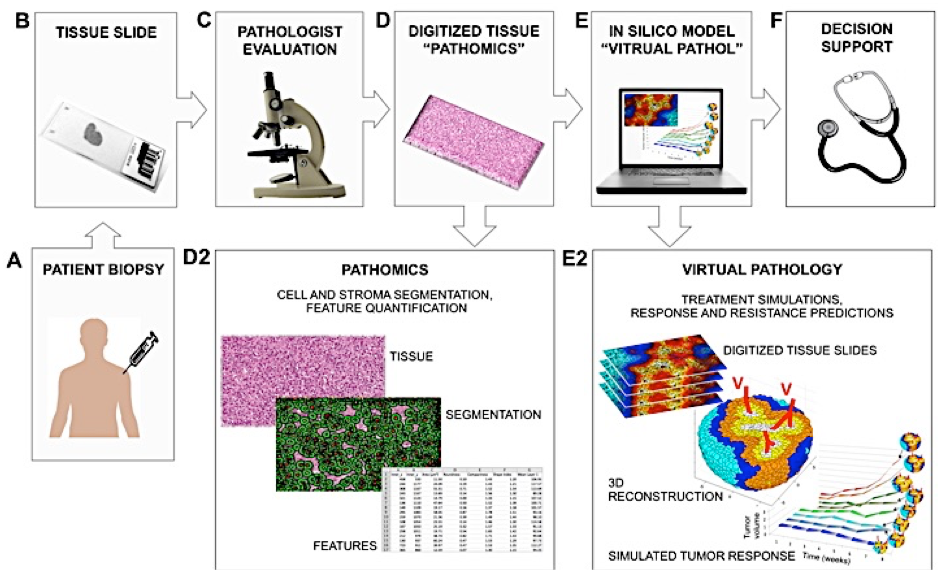
Virtual Clinical Trial “VIRTUOSO” concept for predicting tumor response to chemotherapy based on patients’ biopsy data, quantitative pathology analysis and mathematical modeling [4].

### B. Predicting efficacy of targeted therapy with microPKPD

Targeted therapies are designed to decrease treatment toxicity by selectively aiming at cells that express target receptors. Their efficacy depends on the level of receptors expressed on tumor cells as well as on the physicochemical properties of the tumor tissue that can hinder inratumor drug transport. To predict how to achieve maximal receptor saturation for a given tumor tissue we developed a procedure based on the *microPKPD* model simulations and data from dorsal window chamber (DWC) experiments [7,8]. The schematic of our approach is shown in Figure 2. Images from the DWC experiments (A) were digitized (B), and the tumor tissue architecture was explicitly reproduced in the mathematical model (C). The process of ligand-receptor binding was quantified spatially (D), and the *microPKPD-*simulated drug transport (E) was used for calculations of the association kinetics for various ligand concentrations (F). Exploration of a model parameter space including diffusion, affinity, and release schemes (G) led to scientific predictions (H). Using this approach, we tested which combinations of drug properties (diffusion, affinity), tissue topology (density, cellular loci), drug concentrations, and extravasation rates were critical for optimal drug delivery and desired cellular uptake on an individual cell level.

**Figure 2.**
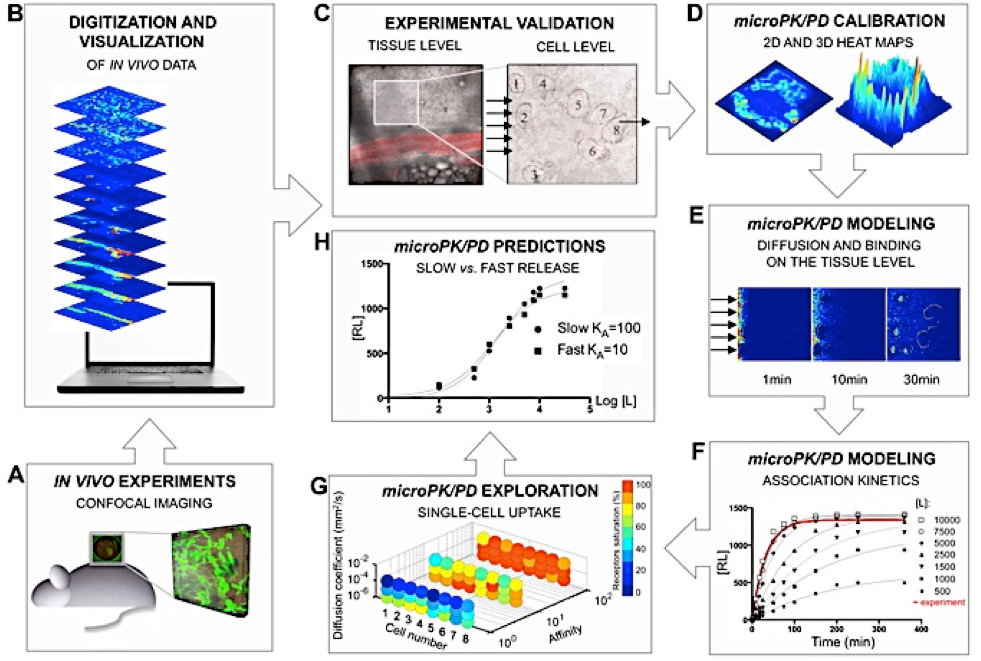
*microPKPD* applied to optimize targeted treatment properties and administration methods to maximize receptor saturation and uptake [7].

### C. Predicting chemotherapy response with Organoid3D

Heterogeneity of tumor microenvironment (mE), and the dynamic metabolic and structural changes that may occur in mE during the treatment may negatively influence efficiency of that therapy. To test interactions between mE and drugs, and to predict how tumors will respond to chemotherapy in various and variable microenvironments, we developed an in silico model of 3D tumor organoids, *Organoid3D.* The model can be calibrated to the properties of a specific cell line and a specific drug, and then used to simulate tumor spheroid growth in a given mE (hypoxia or normoxia, acidity or neutral pH, high or low density of ECM, as well as gradients in all these features), and its response to various drugs and drug schedules. Figure 3 shows model interface, and results of MCF10A-1Ca spheroid growth simulations with no drug (left) and under a low level of doxorubicin (right) in a mild environment (normoxia, neutral pH, low density ECM).

**Figure 3.**
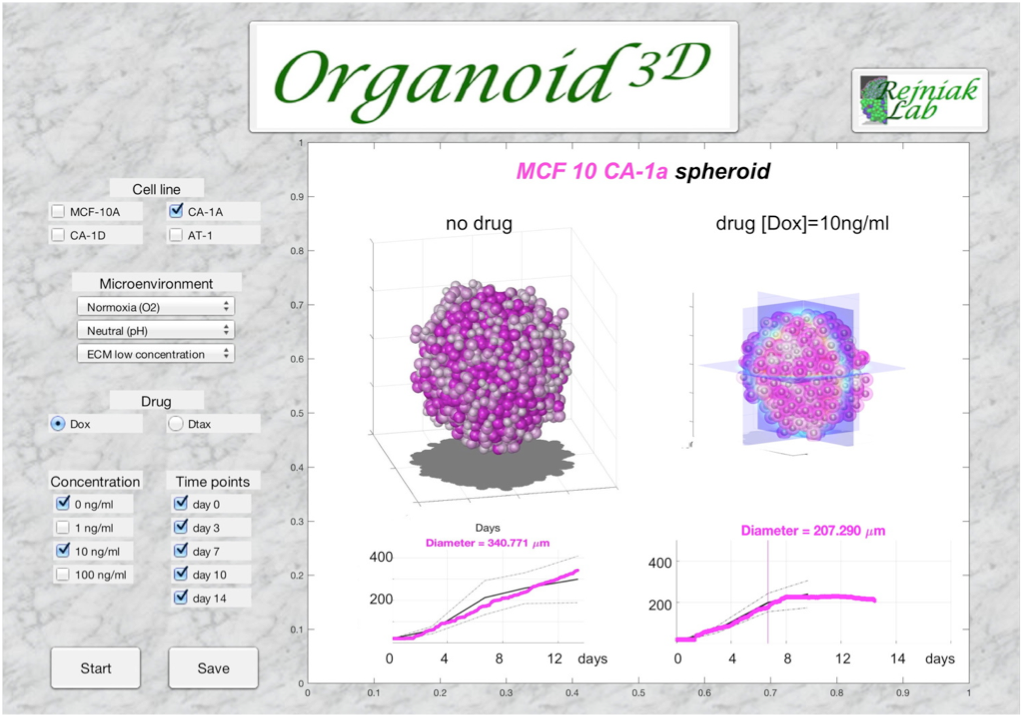
Interface of the in silico model *Organoid3D* for testing tumor response to chemotherapy in various and variable microenvironments [9].

## III. Quick Guide to Mathematical Methods

The *microPKPD* models, including *Organoid3D*, are based on the coupled reaction-diffusion equations of drug and nutrient kinetics [7,8,10], and the particle-spring model for cell mechanics [9,11].

### A. Equations

Drugs are modeled either as discrete particles (Eq.1) that move by a random walk, a discrete diffusion process, and attach to cell membrane receptors according to binding kinetics (Eq.2); or as a continuous density (concentration) of molecules that move by diffusion and advection, and can be internalized by cells (Eq.3). Individual cells are subject to repulsive forces (Eq.4) that arise during cell division; they prevent cells from overlapping and collectively result in passive cell relocation (Eq.5). *microPKPD* is set up in the 2D space, while *Organoid3D* acts in 3D.

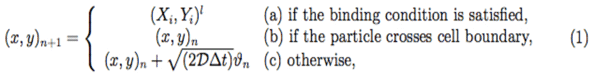

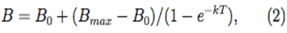

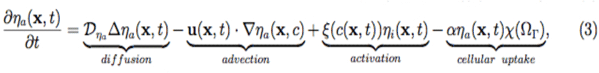

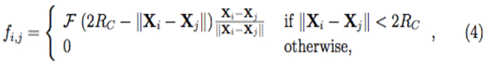

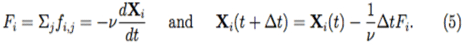

### B. Type of settings in which these methods are useful

The *microPKPD* model can simulate drug transport through the tumor tissue based on either a patient’s or mouse’s histology, as well as drug action on a single-cell level. *Organoid3D* extends this model to 3D space. Both provide useful *micropharmacology* tools for optimizing treatment schedules, testing drug properties to improve their delivery and uptake, as well as predicting how a given tumor will respond to treatments of specific physicochemical properties and whether it will develop resistance.

